# Boosting Felsenstein’s Phylogenetic Bootstrap

**DOI:** 10.1101/154542

**Authors:** F. Lemoine, J.-B. Domelevo Entfellner, E. Wilkinson, T. De Oliveira, O. Gascuel

## Abstract

Felsenstein’s article describing the application of the bootstrap to evolutionary trees, is one of the most cited papers of all time. That statistical method, based on resampling and replications, is used extensively to assess the robustness of phylogenetic inferences. However, increasing numbers of sequences are now available for a wide variety of species, and phylogenies with hundreds or thousands of taxa are becoming routine. In that framework, Felsenstein’s bootstrap tends to yield very low supports, especially on deep branches. We propose a revised version, in which the presence of inferred branches in replications is measured using a gradual “transfer” distance, as opposed to the original version using a binary presence/absence index. The resulting supports are higher, while not inducing falsely supported branches. Our method is applied to large simulation, mammal and HIV datasets, for which it reveals the phylogenetic signal, while Felsenstein’s bootstrap fails to do so.

---

The bootstrap is a widely used statistical method to study the robustness, bias and variability of numerical estimates (*1*). It involves resampling with replacement from the original dataset to obtain replications of the original estimate, and then typically to compute the variance and distribution of this estimate. In 1985, Joseph Felsenstein proposed the application of the bootstrap to assess the robustness (or repeatability) of phylogenetic trees (*2*). Given a sequence alignment and a reference tree inferred on it, the procedure abides by the following steps: (i) resample, with replacement, the sites of the alignment to obtain pseudo-alignments of the same length, (ii) infer pseudo-trees using the very same inference method, and (iii) measure the support of every branch in the reference tree as the proportion of pseudo-trees containing that branch. The usefulness, simplicity and interpretability of this method made it extremely popular in evolutionary studies, to the point that it is generally required for publication of a tree estimate in a wide variety of domains (molecular biology, genomics, systematics, ecology, epidemiology, etc.). Felsenstein’s article was cited more than 32,000 times and is ranked in the top 100 of the most cited scientific papers of all time (*3*). However, the use of Felsenstein’s bootstrap was questioned on biological grounds, notably regarding the assumptions of site independence and homogeneity (*4*). Moreover, the statistical meaning of Felsenstein’s bootstrap proportions (FBP) has been the subject of intense debate (*5*), the main questions being whether FBPs can be seen as the confidence levels of some test, and whether or not they are biased (*6-9*). The mathematical complexity of the tree space and of the related statistical questions (*10*), made all these discussions difficult to follow by most users. The methods to correct the bootstrap support (8, *11-12*) are seldom used, and the original method is still highly cited (~2,000 citations in 2016). Following (*13):* “consensus has been reached among practitioners, if not among statisticians and theoreticians”, and “many systematists have adopted Hillis and Bull’s (6) “70%” value as an indication of support”. The alternatives are the Bayesian posterior probabilities of the tree branches (*14*), which are difficult to obtain with large datasets for computational reasons, and the approximate branch supports (*15-16*), which are fast but provide only a local view. The bootstrap is also computationally heavy, but is easily parallelized and fast algorithms have been designed (*17-18*).

It is commonly acknowledged (*13*) however, that Felsenstein’s bootstrap is not appropriate for large datasets containing hundreds or thousands of taxonomic units (taxa), which are now common thanks to high-throughput sequencing technologies. While such datasets generally contain a lot of phylogenetic information, the bootstrap proportions tend to below, especially when the tree is inferred from a unique gene or a few, as illustrated in Figure 1A with a dataset of ~9,000 HIV-1 group M *DNA polymerase (pol*) sequences. The strongest signal in such a phylogeny generally corresponds to the deep branching of the subtypes. This signal is immediately visible here, and in agreement with the common belief about subtype branching (*19*), but some of the subtypes are not supported (A, B, D, G) and their branching is not supported either (e.g. the grouping of C and H). When using a medium-sized dataset of ~550 randomly selected sequences, the FBPs are higher, with most subtypes supported at 70% or more. However, their deep branching is still unresolved (Fig. S8).

**Figure 1:**
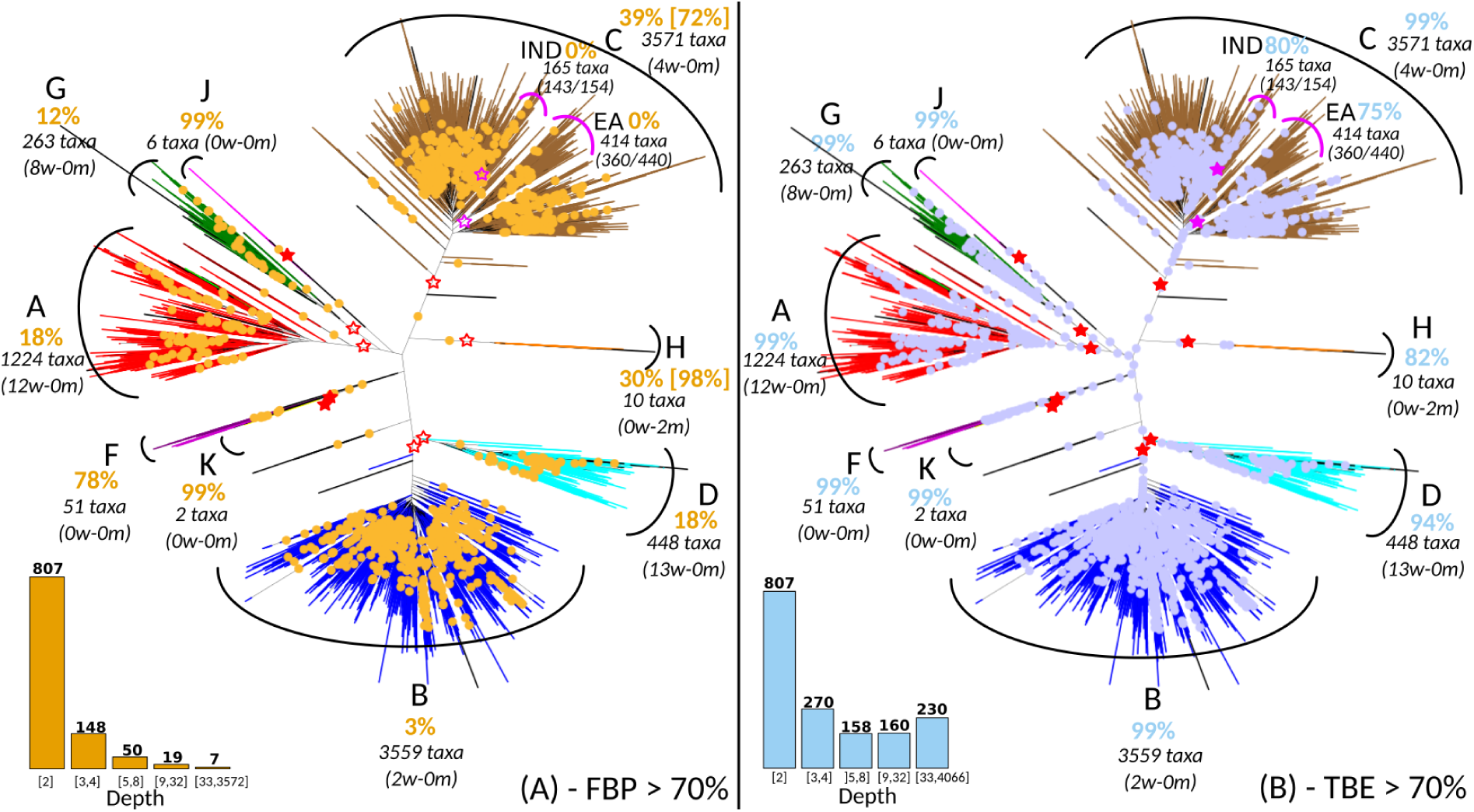
Felsenstein (FBP) and transfer (TBE) bootstrap supports on the same tree with 9,147 HIV-1M *pol* sequences. Subtypes are colorized; recombinant sequences are black; dots correspond to branches with support >70% (following (*6*)). Supports are given for the tree clades that are closer to the subtypes (red stars, filled when support >70%); for each of these clades we provide the number of wrong (*w*) taxa that do not belong to the corresponding subtype, and the number of missing (*m*) taxa that belong to the subtype but not to the clade. For the C and the H, these clades are not supported by FBP but there exist neighboring clades with FBP >70%, and these are shown in brackets. The same approach is applied to the C sub-epidemics in India (IND) and Eastern Africa (EA); the ratio provides the coverage of the clade, i.e. the number of studied (e.g. Indian) taxa in the clade versus the total number of those taxa in the dataset. The South American clade (SA, not shown, included in EA) is supported by TBE but not by FBP (73% vs. 14%, 15 taxa, 14/14). The histograms provide the number of branches with support >70% depending on branch depth, which is measured by the number of taxa in the smaller of the two clades defined by the given branch. (Sup. Text 3)

The reason for such degradation is explained by the core methodology of Felsenstein’s bootstrap. A replicated branch must match a reference branch exactly to be accounted for in the FBP value. A difference of just one taxon (highly likely with large datasets) is sufficient for the replicated branch to be counted absent while it is nearly identical to the reference branch (*13*, *20*). There are many biological and computational reasons for the existence of “rogue” taxa with unstable phylogenetic positions: convergence, recombination, sequence and tree errors, etc. The standard approach (*20-24*) is to remove these taxa and relaunch the analysis, but this is statistically questionable and computationally expensive.

Our approach has a simple but sound statistical basis. We replace the branch presence proportion (*i.e*. the expectation of a {0,1} indicator function) of Felsenstein’s bootstrap, by the expectation of a refined, gradual function in the [0,1] range, quantifying the branch presence in the bootstrap trees. We use the “transfer” distance (*25-27*), where the distance δ(*b*,*b*\*) between a branch *b* of the reference tree *T* and a branch *b\** of a bootstrap tree *T\** is equal to the number of taxa that must be transferred (or removed), in order to make both branches identical (*i.e*. both split identically the set of taxa). To measure the presence of *b* in *T\**, we search the branch in *T\** that is closest to *b* and use the “transfer index” ϕ(*b*, *T* *) = *Min*_*b*\*∈*T*\*_ {δ( *b*, *b*\*)}. This index has several important and useful properties. Let *l* be the number of taxa. Any branch *b* splits the taxa into two subsets; let *p* be the size of the smaller subset induced by *b* (*p* ≤ *l* − *p*). We have (Sup. Text 1):

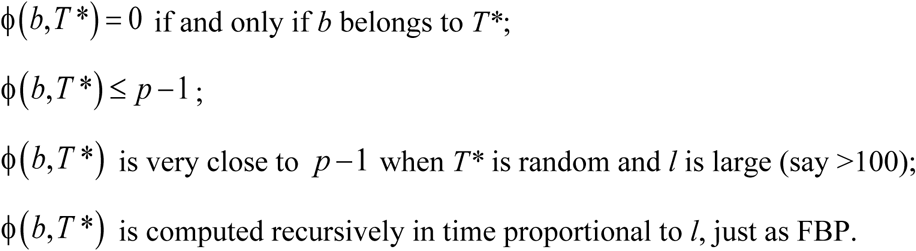

Based on these properties, we define the transfer bootstrap expectation (TBE) as:

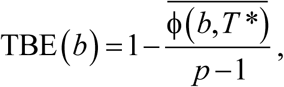

where the numerator is the average transfer index among all bootstrap trees. It is easily seen that TBE ranges from 0 to 1, where 0 means that the bootstrap trees are random regarding *b*, and 1 means that *b* appears in all bootstrap trees. Moreover, considering the same set of bootstrap trees, TBE(*b*) is necessarily larger than FBP(*b*) and the difference is substantial for deep branches, while TBE(*b*) = FBP(*b*) when *b* defines a (shallow) “cherry” (i.e. *p =* 2). Importantly, TBE does not support wrong branches and induces the same, low amount of false positive errors as FBP when used with common thresholds, typically 70% (6) or higher (Fig. 2C-D, S3-S4, S10-S11).

**Fig. 2:**
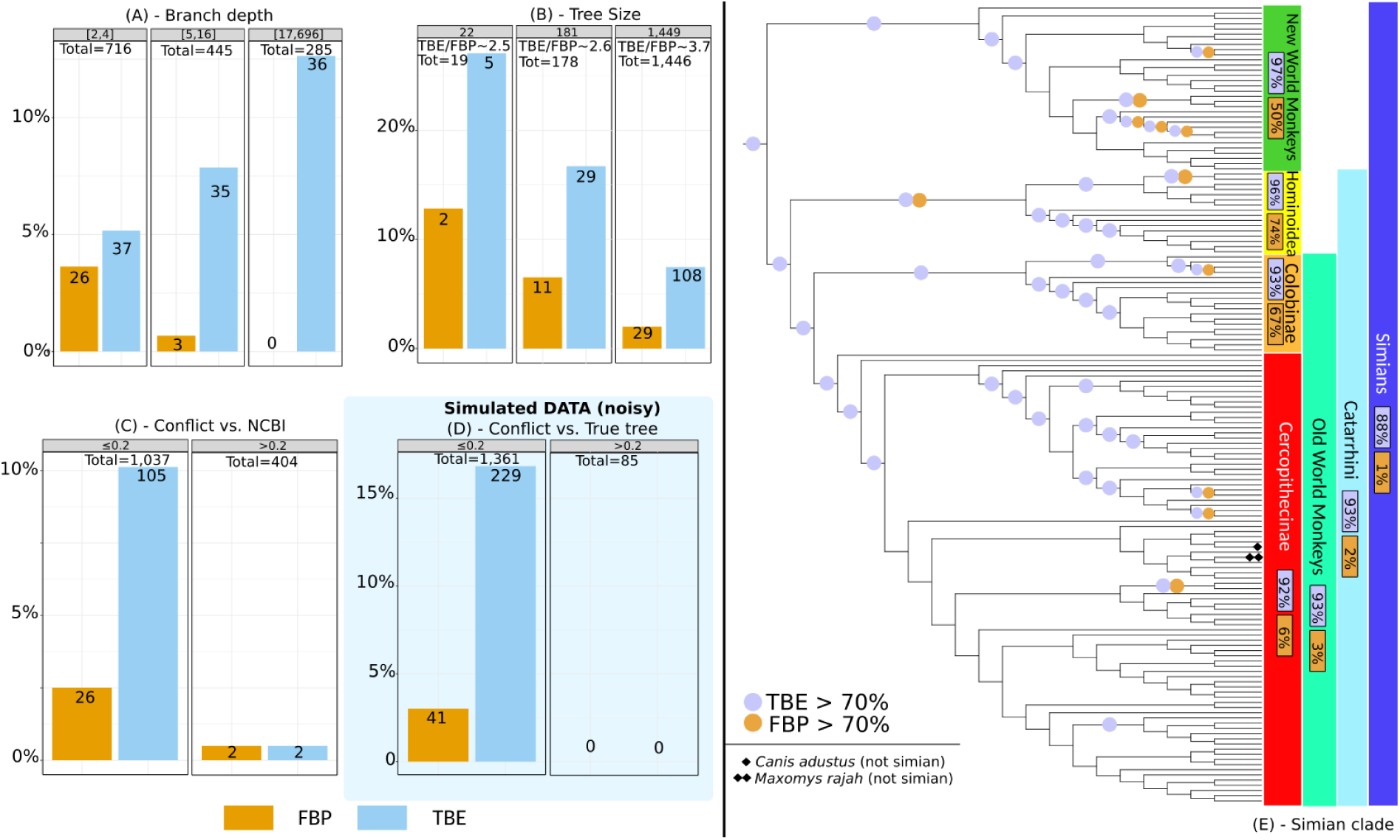
Felsenstein (FBP) and transfer (TBE) bootstrap supports with 1,449 COI-5P mammal sequences – FastTree phylogeny. Graphs A to D refer to branches with supports >70%, with the vertical axis denoting the percentage of these branches in a given condition (e.g. (B): 19 internal branches with 22-taxon trees, and 2/19 ≈ 10% of branches with FBP >70%). (A) Supports regarding branch depth (see note to Fig. 1). (B) Supports regarding tree size (*i.e*. number of taxa). (C) Supports regarding percentage of quartet conflicts with NCBI taxonomy. (D) Same as (C) but regarding the true tree used for simulations. (E) Results with the simian clade, all simian sequences are included, but two additional non-simian sequences are added, one rogue taxon (*Maxomys rajah*) and one stable but erroneous taxon with partial sequence (*Canis adustus)*. (Sup. Text 2)

All these properties are highly desirable: easy computation, higher support than FBP, low false positive rate. Moreover, TBE has a simple and natural interpretation; for instance, with *l =* 1,000 and *p =* 200, TBE(*b*) = 95% means that on average (200 − 1)×0.05 ≈ 10 taxa have to be transferred to recover *b* in a bootstrap tree, while 990 taxa are stable regarding *b*. Lastly, we can define an instability score for each taxon, based on the number of times it is transferred in TBE computations.

We first studied the advantage of TBE on a large phylogeny of 1,449 mammals, obtained from a usual barcoding marker (COI-5P; Sup. Text 2). The reference and bootstrap trees were inferred by maximum likelihood from the protein alignment (527 sites) using both FastTree (*28*) and RAxML with rapid bootstrap (*17*) to check that similar conclusions were drawn with different inference methods. To study the impact of the number of taxa, we randomly selected small (22 taxa) and medium (181 taxa) datasets and performed the same analyses. The results were compared to the NCBI taxonomy (*29*), which represents current thinking about mammals’ evolutionary history, with unresolved nodes in case of uncertainty. Results indicate clearly that TBE provides some support for deep branches, while FBP does not (Fig. 2A, S3-S4). As expected, the supports of shallow branches are similar, and the impact of TBE is more pronounced with a large number of taxa, but still of interest with medium-sized datasets (Fig. 2B, S3-S4). Obviously, getting higher supports only makes sense if the number of falsely supported branches remains low. For all inferred branches, we thus measured the quartet-based percentage of contradictions with the NCBI taxonomy. TBE supports a larger number of weakly contradicted branches than FBP, which fulfills one of its objectives (nearly correct branches must be supported), while the number of highly contradicted branches remains extremely low and nearly identical for both approaches (Fig. 2C, S3-S4), a result confirmed by simulations (Fig. 2D, S10-S11). The advantage of TBE appears clearly when inspecting the tree clades. For example (Fig. 2E and S5 for RAxML analysis), the simian clade has a strong support with TBE, while FBP is nearly null due to a high number of rogue taxa in the bootstrap trees, and the same holds for several sub-clades. Similar results (Fig. S6) were found with other well-established clades (cetaceans, marsupials, monotremes…), and using RAxML. In comparison, the removal of rogue taxa (*24*) does not significantly improve FBP (8 and 3 taxa are removed with FastTree and RAxML, respectively, but the number of branches with FBP >70% remains the same).

Next, we applied our method to a large dataset of 9,147 HIV-1 group M *pol* sequences (Sup. Text 3). Such large datasets are increasingly common in molecular epidemiology and phylodynamics (*31*). We only retained sequences annotated as non-recombinant by the Los Alamos HIV-1 DB using a fast filtering approach. Among these, 48 recombinant sequences were still detected by jpHMM (*30*). These 48 sequences were kept in the analyses to study the impact of recombinant sequences, as their presence is inevitable in any HIV dataset. As opposed to mammals, the tree topology of HIV-1M strains is essentially unknown. Moreover, it is intrinsically unstable as reconstructing a tree with so many relatively short and possibly recombinant sequences is a very hard task. Thus, the main expectation is to observe a clear separation between the subtypes. We built the reference and bootstrap trees using FastTree on the DNA sequence alignment (1,043 sites), and performed the same analyses using smaller subsets of 35 and 571 sequences. While the deep branching of the subtypes (*19*) is poorly supported by FBP (Fig. 1A), it becomes apparent with TBE, where all subtypes have a support larger than 80%, and close to 100% in most cases (Fig. 1B, S8). The same holds true with other well described clades; for example, TBE supports the identification of regional variants of HIV-1 subtypes that are of epidemiological importance: the East-African, Indian, and South American subtype C variants, which FBP fails to support. TBE provides a substantial support to a much larger number of deep branches. Again, the advantage in using TBE is higher with large data sets (Fig. S7), but still apparent with 571-taxon datasets, where the deep subtype branching and C sub-epidemics are supported by TBE but not FBP (Fig. S8). An important feature of TBE is that the supports may be non-local, but attached to “caterpillar-like” paths (Fig. 1B, e.g. subtype C). With HIV-1M data, this corresponds to the fact that the subtype roots are usually not well defined due to recombinant and ancient sequences. Moreover, the instability score among recombinant sequences is clearly higher than in the sequences not detected as recombinant (Fig. S9), supporting the biological soundness of the approach and its power in detecting recombination and rogue taxa.

Lastly, to check that TBE does not support erroneous branches, we performed extensive computer simulations with various tree sizes and phylogenetic signal levels (Sup. Text 4). We also added unstable taxa bearing a weaker phylogenetic signal than the others. The results are highly similar to those with real data regarding the support of deep branches and the tree size (Fig. 2A-B, S10-11). In all the conditions we examined, the false positive rate was very low and nearly identical to that obtained with FBP, and the rogue taxa exhibited lower stability (Fig. S12).

The transfer bootstrap thus provides a measure of branch robustness. The results clearly demonstrate its usefulness, especially with deep branches and large datasets, where branches known to be essentially correct are supported by TBE but not by FBP. Though higher, these supports are not impaired with falsely supported branches. Importantly, they are easily interpreted as fractions of taxa to be transferred to recover the inferred branches. It is thus likely that future users will choose the support level depending on the phylogenetic question being addressed (e.g. lower TBE support threshold with HIV and possibly recombinant sequences, than with mammals), rather than on the basis of empirical rules such as the common 70% level used with FBP. Moreover, our experiments demonstrate the ability of the transfer index in detecting unstable taxa responsible for low supports. Lastly, the approach is applicable to rapid bootstrap ((*17-18*), Fig. S4, S6) and could be extended to Bayesian branch supports.

## Acknowledgements

We thank Frederic Delsuc, Susan Holmes and Leonid Chindelevitch for help and suggestions. This work was supported by the EU-H2020 Virogenesis project (grant number 634650 - EW, TDO, and OG) and by the H3ABioNET project (NIH grant number U41HG006941 - JBDE).

## Data and software

All our data and software are available from Booster web site and GitHub (http://booster.c3bi.pasteur.fr ; https://github.com/evolbioinfo/booster

## Author contributions

OG designed the research; FL, JBDE and OG performed the research; FL and JBDE implemented the algorithms; FL realized the website and GitHub repositories; FL performed the analyses and graphics, with the help of EW and TDO for HIV; OG wrote the paper, with the help of all coauthors.

## Supporting Online Material

(Sup. Text 1): Methods and software, Fig. S1-S2

(Sup Text 2): Mammals dataset and analyses, Fig. S3-S6

(Sup Text 3): HIV dataset and analyses, Fig. S7-S9

(Sup Text 4): Simulated datasets and analyses, Fig. S10-S12

## Methods (Supporting Text 1)

### Transfer distance and index: definitions and properties

The transfer distance (*25*), also called R-distance (*26*), was introduced to compare partitions in cluster analysis. In this context, the transfer distance is equal to the minimum number of elements to be transferred (or removed) to transform one partition into the other. Tree branches are commonly seen as bipartitions or splits, as a branch divides the taxa into two subsets situated on its two sides. The most used topological distance between two trees is the Robinson and Foulds distance (*32*), which is equal to the number of bipartitions that belong to one tree but not the other. As quoted by (*27*), the bipartition distance is overly sensitive to some small tree changes, possibly involving a unique taxon. Those authors proposed using the transfer distance and designed algorithms to compute a more robust “matching” distance between trees, a different task, but related to the aim of this article. In the following, we first provide basic definitions (see (*33*) for a text book on phylogenetic trees), and then demonstrate the properties of the transfer distance in a bootstrap context.

Let X be a fixed set of *l* taxa. An X-tree is a phylogenetic tree with *l* leaves labeled by the taxa of X. All (reference, bootstrap) trees discussed here are X-trees, meaning that they bear on the same set of *l* taxa. Any branch of an X-tree defines a bipartition of X, and the topology of an X-tree can be recovered from its bipartition set. Thus, we will use both terms (branch, bipartition) indifferently, depending of the context. Any bipartition *b* of X can be encoded by a {0,1} vector **v**( *b*) of length *l*, where the taxa on the same side of the bipartition are encoded by the same value. Note that *b* is also encoded by 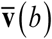 the negation of **v**(*b*) (i.e. the 0s are turned into 1s and vice versa). Moreover, the smaller of the two subsets induced by a bipartition *b* will be called here the “light side” of b, and *p* will denote the size of the light side of *b* (*p* ≤ *l* − *p*). One says that a bipartition is “trivial” when it has a unique taxon in its light side (*p* = 1). An X-tree defines *l* trivial bipartitions corresponding to each of the taxa. These trivial bipartitions are contained in every X-tree, while the other non-trivial bipartitions define the core of the tree topology and are the central subject of phylogenetic studies.

The transfer distance δ(*b*, *b*\*) between a bipartition *b* of the reference tree *T* and a bipartition *b\** of a bootstrap tree *T\** is equal to the number of taxa that must be transferred (or removed) to make both bipartitions identical. The transfer distance is easily defined and computed using the Hamming distance *H* between **v**(*b*) and **v**(*b*\*) :

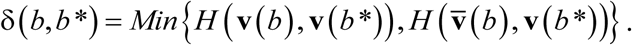

To measure the presence of *b* in *T*\*, we search the bipartition in *T\** that is closest to *b* and use the transfer index ϕ(*b*, *T* *) = *Min_b_*_*∈*T*\*_ {δ(*b*, *b* *)}. Based on above definitions, δ(*b,b*\*) = 0 if and only if **v**(*b*) and **v**(*b*\*) define the same bipartition of X. Thus, the transfer index satisfies:

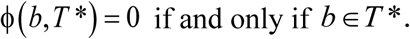

Moreover, let *b* be any given bipartition of *T* and *t* be a taxon in the light side of *b*. The trivial bipartition *b*\* *=* {*t*}/X –{*t*} is found in any bootstrap tree *T\** and δ(*b*, *b*\*) = *p* −1. There may well be another bipartition closer to *b* in *T\**, but at least this ensures that:

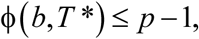

and thus, the transfer support *TS* satisfies:

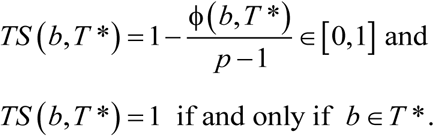

Let 1*_b_*(*T* *) be the indicator function equal to 1 when *b* ∈ *T* * and 0 otherwise. For any bipartition *b* and tree *T\**, we have 1*_b_*(*T*\*) ≤ *TS*(*b, T*\*). The Felsenstein bootstrap proportion (FBP) is equal to the average of 1*_b_*(*T*\*) over the set of bootstrap trees, while the transfer bootstrap expectation (TBE) is equal to the average of *TS*(*b*, *T*\*). Thus, when using the same set of bootstrap trees, we necessarily have FBP(*b*) ≤ TBE(*b*). When *b* is a “cherry” (i.e. *p* = 2), we have 1*_b_*(*T*\*) = *TS*(*b*, *T* *) and thus FBP(*b*) = TBE(*b*). With deeper bipartitions, we generally observe that in the presence of a significant phylogenetic signal, only a small number of taxa need to be transferred to make *b* identical to a bipartition in *T\**, while the strict presence of *b* in *T\** is relatively rare; the difference between FBP(*b*) and TBE(*b*) is then substantial.

### Recursive computation of the transfer index

(*27*) describes a recursive algorithm to compute all transfer distances between any given bipartition *b* of *T* and all bipartitions of another tree *T’*. This algorithm is easily transformed to compute the transfer index.

The principle is as follows:

(1) Map all the leaves of the light side of *b* to 0, the others to 1, and apply the same mapping to the leaves of *T\**. Moreover, root *T\** at any internal node.
(2) With a single postorder tree traversal, one can compute the number of leaves labeled 0 and the number of leaves labeled 1 for every subtree in *T\**.
(3) Let *l*_0_ be the number of leaves labeled 0 and *l*_1_ be the number of leaves labeled 1 in the subtree attached below a given bipartition *b\**. The transfer distance between *b* and *b\** is given by (think to the missing 0s and the 1s to be removed in *b\** subtree, and vice versa):

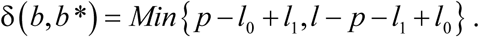 This distance can be computed during the postorder traversal as well as the transfer index ϕ(*b*, *T* *), which is the minimum of δ(*b*, *b*\*) for all bipartitions of *T\**.

This algorithm has linear time complexity, and thus computing TBE for all bipartitions in *T* with *r* bootstrap replicates has a time complexity in *O*(*rl*^2^). FBP has the same time complexity, but very efficient implementations have been developed (*e.g*. using bit vectors to encode bipartitions). In practice, computing all TBE supports with 4,000 taxa and 1,000 replicates requires less than one hour (5 core Intel Xeon 3.5GHz), which is negligible compared to the time required to infer the reference and bootstrap trees.

### Expected transfer index with random trees and TBE distribution

We have seen that the transfer index satisfies ϕ(*b*, *T*\*) ≤ *p* − 1. We show in this subsection that the expected transfer index is very close to this upper-bound with random “bootstrap” trees, when the number of taxa is large enough. Consequently, the transfer bootstrap expectation of any branch 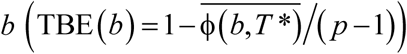 is close to 0 when the bootstrap trees seem to be random and do not contain any signal regarding *b*. This property explains why moderate supports, for example 70% as used throughout the article, are sufficient to reject poor branches, as a branch support of 70% cannot be observed by chance.

We first provide a simple argument to explain this result, based on the expected transfer distance between a fixed bipartition *b* and a random bipartition *b\** with fixed light-side size *p*\*. Let *x* = *p/l* denote the proportion of taxa in the light side of *b* ( *x* ≤ 1 − *x* since *p* ≤ *l* − *p*). Both bipartitions *b* (fixed) and *b\** (random) define four taxon subsets, the expected sizes of which are given by the following contingency table (each cell follows a hypergeometric distribution):

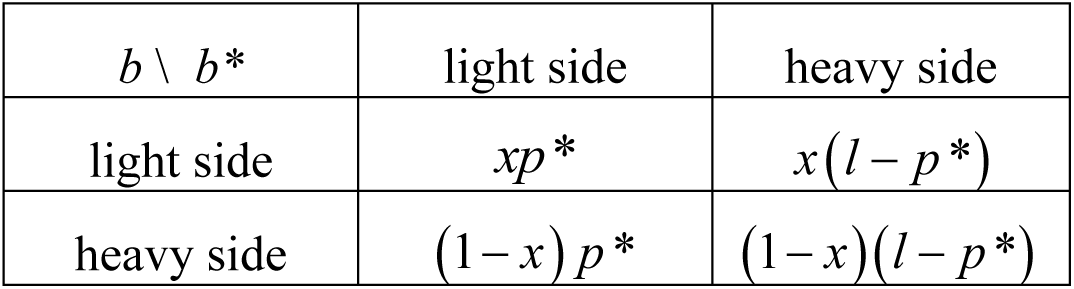

It is easily seen that under above assumptions, the expected transfer distance between *b* and *b\** is equal to the sum of the anti-diagonal terms, that is:

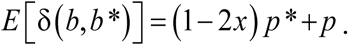

Since p* *>* 0 and (1 − 2*x*) ≥ 0, we have:

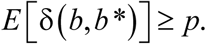

This result shows that the expected transfer distance between *b* and *b\** is larger or equal to *p*, for any value of *p* and *p\**. Moreover, the lower *p\**, the closer the expected transfer distance is top. As a first approximation, we thus see that the transfer index should be close to its upper-bound *p* − 1, since it is equal to the minimum of distances which taken separately are all expected to be larger than *p*.

However, these distances fluctuate around their expected values, and their minimum may be lower than the minimum of their individual expectations, especially with small samples (i.e. low number of taxa). We performed computer simulations to measure the extent of this phenomenon and the validity of the *E* [ϕ(*b*, *T*\*)] ≈ *p* − 1 approximation. We used four tree sizes: *l* = 16, 128, 1,024 and 8,192 taxa, and four models of random phylogenetic trees: caterpillars (fully imbalanced), PDA, Yule-Harding, and perfectly balanced (*33*). For the bipartition b, all possible integer values of *p* in the [2, *l*/2] range were used. The number of random “bootstrap” trees was equal to 1,000, and we performed 100 runs per tree size.

Results are displayed in Fig. S1-S2. With *l* ≥ 1,024, the average transfer index with random trees is surprisingly close to the upper bound *p* − 1, and the approximation is already satisfying with *l =* 128. Moreover, the results are nearly the same for the four random tree models, suggesting that the property holds in a number of settings. As expected, the approximation is better with small *p*. Indeed, remember that the upper bound *p* − 1 is obtained with a trivial bipartition *b\** made of a unique taxon belonging to the light side of b. When a cherry in *T\** contains two taxa from the light side of b, then ϕ(*b,T* *) ≤ *p* − 2 . Similar deviations are observed with subtrees in *T\** containing a large fraction of taxa belonging to the light side of b. The largerp, the higher the probability for such event to occur. Note, however, that large values of *p* (i.e. *p* ≈ *1/*2) are relatively rare for most tree models (*e.g*. Yule-Harding). Looking at the distribution of TBE, we see that having TBE larger than a moderate threshold (say 50%) is very unlikely, even with 16 taxa, thus explaining that TBE rarely supports poor branches with real and simulated data (Fig. 2C-D, S3-S4, S10-S11).

**Fig. S1:**
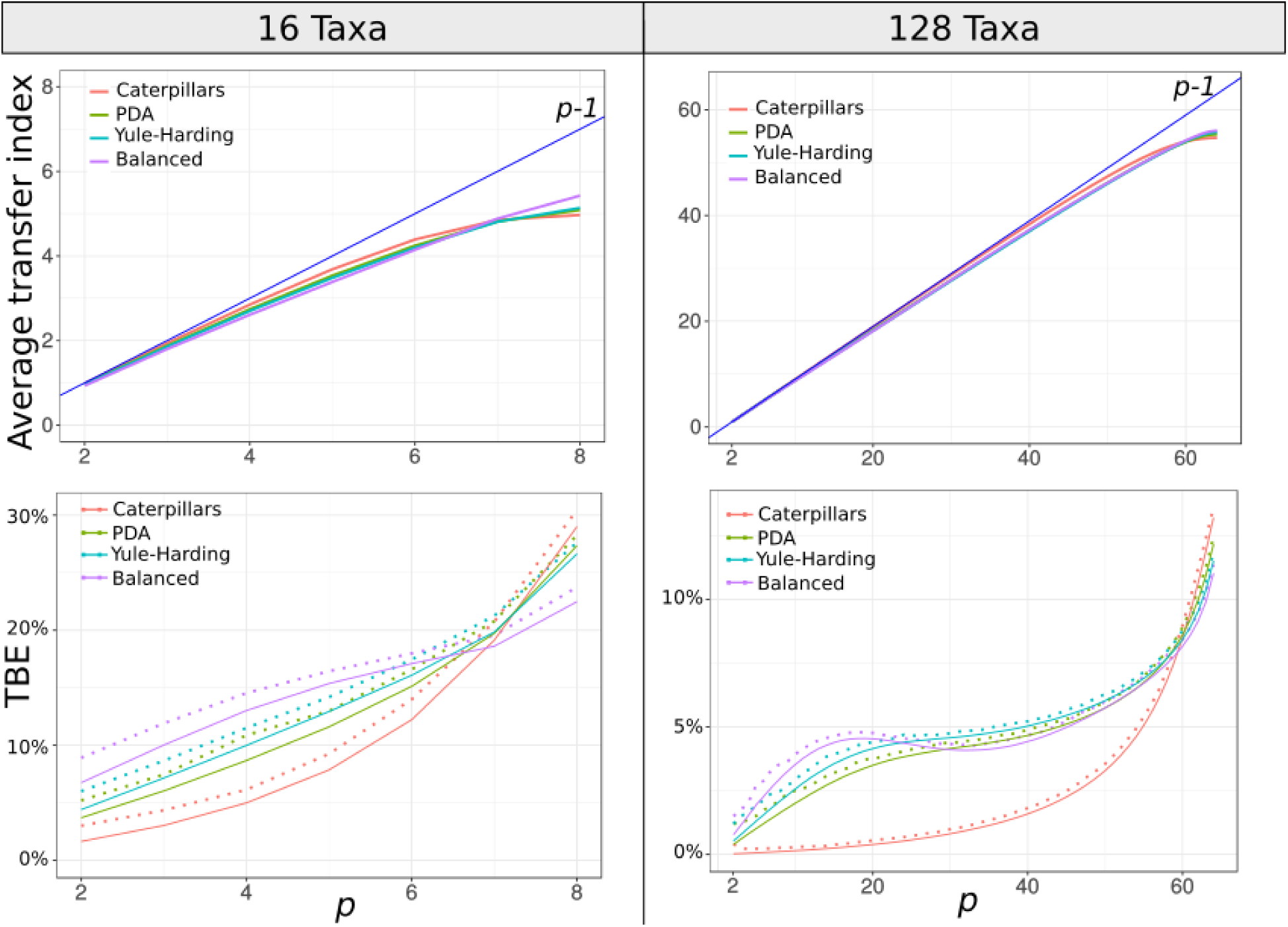
Transfer index and TBE support expectation with random trees of 16 and 128 taxa. For each number of taxa and random tree model, we compare the transfer index average over 100 runs with upper-bound *p*-1 (top graphs). We also compare the average transfer bootstrap support (TBE) to 0, and provide (dashed lines) the maximum value observed among 100 runs, thus approximating the 1% quantile of the distribution (bottom graphs).

**Fig. S2:**
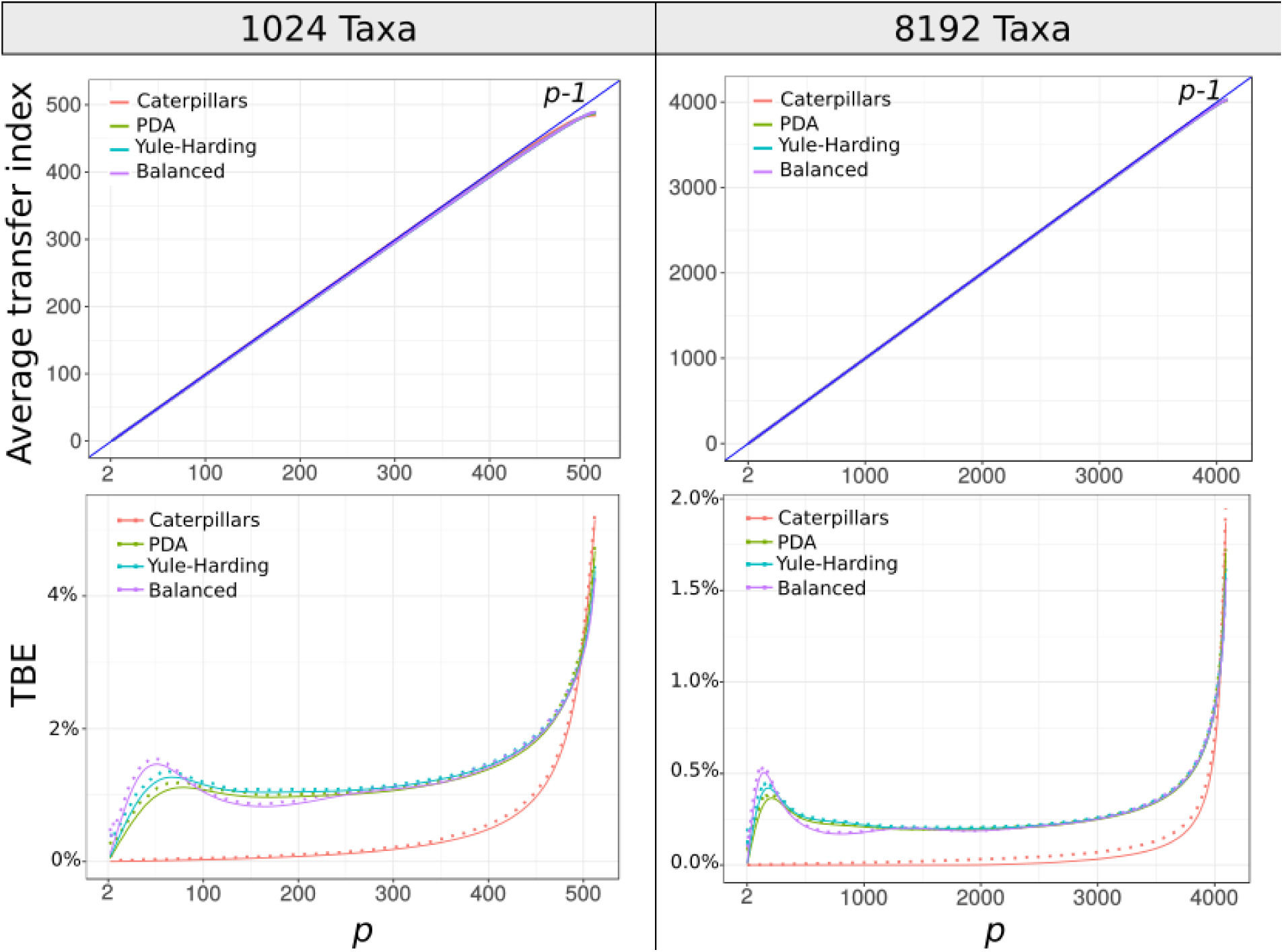
Transfer index and support expectation with random trees of 1,024 and 8,192 taxa. See note to Fig. S1.

### Software programs, web server

We developed several tools to compute the transfer bootstrap. We first implemented a command line tool in C, “Booster”, available at https://github.com/evolbioinfo/booster. This tool can compute TBE as well as FBP supports. It takes two files as input: (1) a reference tree file in Newick format, and (2) a bootstrap tree file in Newick format, containing all bootstrap trees. A number of software programs can be used to infer trees from MSAs and produce these reference and bootstrap files in the desired format, such as RAxML, FastTree and PhyML, as used in this article, and many others (see examples in Booster GitHub repository).

We also developed “BoosterWeb” (http://booster.c3bi.pasteur.fr), a freely available web interface, which allows users to compute bootstrap supports (TBE and FBP) easily without installing any tool on their own computer. Computations are launched on the Institut Pasteur cluster throughout a Galaxy instance. As for the command line tool, two input files are needed: (1) a reference tree file in Newick format, and (2) a bootstrap tree file in Newick format. We propose a basic visualization of the resulting tree highlighting highly supported branches at a given threshold. The resulting tree can be uploaded in one-click on iTOL ((*34*); http://itol.embl.de/) for further manipulation. Moreover, BoosterWeb is self-contained and can be easily installed on any desktop computer (Windows, MacOS, and Linux) by downloading the BoosterWeb executable.

For the sake of reproducibility, all the analyses described in this article were implemented in the NextFlow workflow manager ((35); https://www.nextflow.io/), and are accessible at https://github.com/evolbioinfo/booster-workflows. All the tools we used are publicly accessible and listed with their version on this repository. All our data are available for download (mammals COI-5P alignment, HIV-1 *pol* alignment, simulated data). The software programs that we developed to manipulate data, change formats, etc., are available for download at http://github.com/fredericlemoine/goalign and http://github.com/fredericlemoine/gotree for trimming alignments and trees.

## Mammal dataset and analyses (Supporting Text 2)

### Data, alignments, tree building, comparisons with NCBI

We downloaded all aligned mammals COI-5P amino-acid sequences from the Barcode of Life Data System (BOLD - http://www.barcodinglife.org - date of access: September 2015). We removed all sequences shown to be identical among several species, kept one sequence per species (several gene versions are available for some species, but no paralogs), and converted the resulting multiple alignment (1,449 sequences, 527 sites) into Fasta format. This alignment was subsampled to study the impact of tree size. We randomly drew 8 samples with 1/8th of the sequences (i.e. 181) and 64 samples with 1/64th of the sequences (i.e. 22). We then generated 1,000 bootstrap alignments for the full alignment and each of the 72 sub-sampled alignments by drawing sites with replacement.

We used FastTree ((*28*); options: –nopr –nosupport –wag –gamma) to infer trees from each of these reference (1+8+64=73) and bootstrap (73,000) alignments. To ensure that the results and conclusions were independent of the tree building method, we also performed the same analyses using RAxML with rapid bootstrap ((*17*); options –f a –m PROTGAMMA –c 6 –T 10 –p $RANDOM –x $RANDOM -#1000). The FBP and TBE supports for the (73 × 2) reference trees were computed using Booster (command-line C version). All trees were drawn using iTOL and are available on Booster GitHub repository, along with the sequence alignments. To see if rogue taxa removal improves FBP supports, we ran RAxML rogue-detection tool ((*12*); options –J MR_DROP –z bootstrap_trees –m PROTGAMMAWAG -c 6 -T 4) and recomputed FBP supports without selected taxa.

The FastTree and RAxML complete tree topologies were compared to the NCBI taxonomy (https://www.ncbi.nlm.nih.gov/taxonomy), which was converted to Newick format and reduced to the 1,444 taxa common to both our alignment and the NCBI taxonomy. This NCBI tree is not fully resolved and summarizes common belief about the evolutionary history of mammals, resulting from a number of phylogenetic studies based on numerous markers. The unresolved part of the NCBI tree (~4.35 descendants per node on average, instead of 2 for a fully-resolved tree) corresponds to the unknown or uncertain part of that history. To cope with irresolutions, we used quartets to compare the (fully-resolved) inferred trees to the NCBI tree. A quartet is a tree topology with 4 taxa; AB|CD is the standard notation for quartets, indicating that taxa A and B form a cherry separated by an internal branch from the cherry formed by C and D; a quartet is unresolved when the 4 taxa are connected to a single central node. A bipartition *b* induces a quartet AB|CD when A and B belong to the same side of *b*, and C and D to the other side. We used tqDist (*36*) to count the number of quartets induced by the reference branches, which appeared to be contradictory with the quartets induced by the NCBI tree and its bipartitions; for example, AB|CD was found in the studied branch, while AC|BD was found in the NCBI tree. Unresolved quartets of the NCBI tree were not counted as contradictory, as they are just unknown and the inferred resolution could be correct. The number of contradicted quartets was divided by the total number of quartets induced by the studied branch, to obtain a normalized measurement in the [0, 1] range (0: no contradiction; 1: all induced quartets are contradicted). We used the same approach to check the accuracy of the FastTree and RAxML tree topologies, comparing the whole set of quartets induced by the inferred tree to those induced by the NCBI tree.

### Overall comparison of FBP and TBE (Fig. 2, S3-S4)

- The RAxML topology is clearly closer to the NCBI taxonomy than the FastTree topology is (27% versus 38% of contradicted quartets, respectively). However, the RAxML topology is still relatively poor, as expected in this type of phylogenetic study based on a unique marker (see Fig. S6 and further discussion). Despite this difficulty, FBP and TBE perform well with both programs as they give supports larger than 70% to a very low number of contradicted (>20%) branches.
- The supports obtained with RAxML are higher than FastTree’s (47 versus 29 branches with FBP>70%, and 158 versus 108 with TBE>70%, for RAxML and FastTree, respectively). Part of the explanation could be that the RAxML tree is more accurate than that of FastTree, and thus better supported. Another factor is that the rapid bootstrap tends to be more supportive than the standard procedure (e.g. (*16*)). Indeed, the rapid bootstrap uses already inferred trees to initiate tree searching, and thus tends to produce less diverse bootstrap trees than the standard (slower) procedure, which restarts tree searching from the very beginning for each replicate.
- Despite these differences between FastTree and RAxML with rapid bootstrap, similar conclusions are drawn when comparing FBP and TBE: FBP supports very few deep branches, while TBE supports a larger number of them; TBE is especially useful with large trees; both methods support a very low number of contradicted branches.
- Comparing the three cutoffs, we see that with 50% the selected branches are still weakly contradicted, especially with FBP; with TBE, as expected, the fraction of contradicted branches (>20%) is a bit higher but still low (~5%). With 90%, very few branches are selected (~7% with TBE), thus justifying the use of 70% for TBE as is standard with FBP. However, it is likely that, rather than relying on such arbitrary value, most users will choose cutoffs based on the interpretation of TBE and the specificity of their phylogenetic analysis, and typically argue that a given branch is well supported since only a few taxa need to be transferred and are unstable. Moreover, these taxa will possibly be identified thanks to the instability score (Fig. S9, S12).

**Fig. S3:**
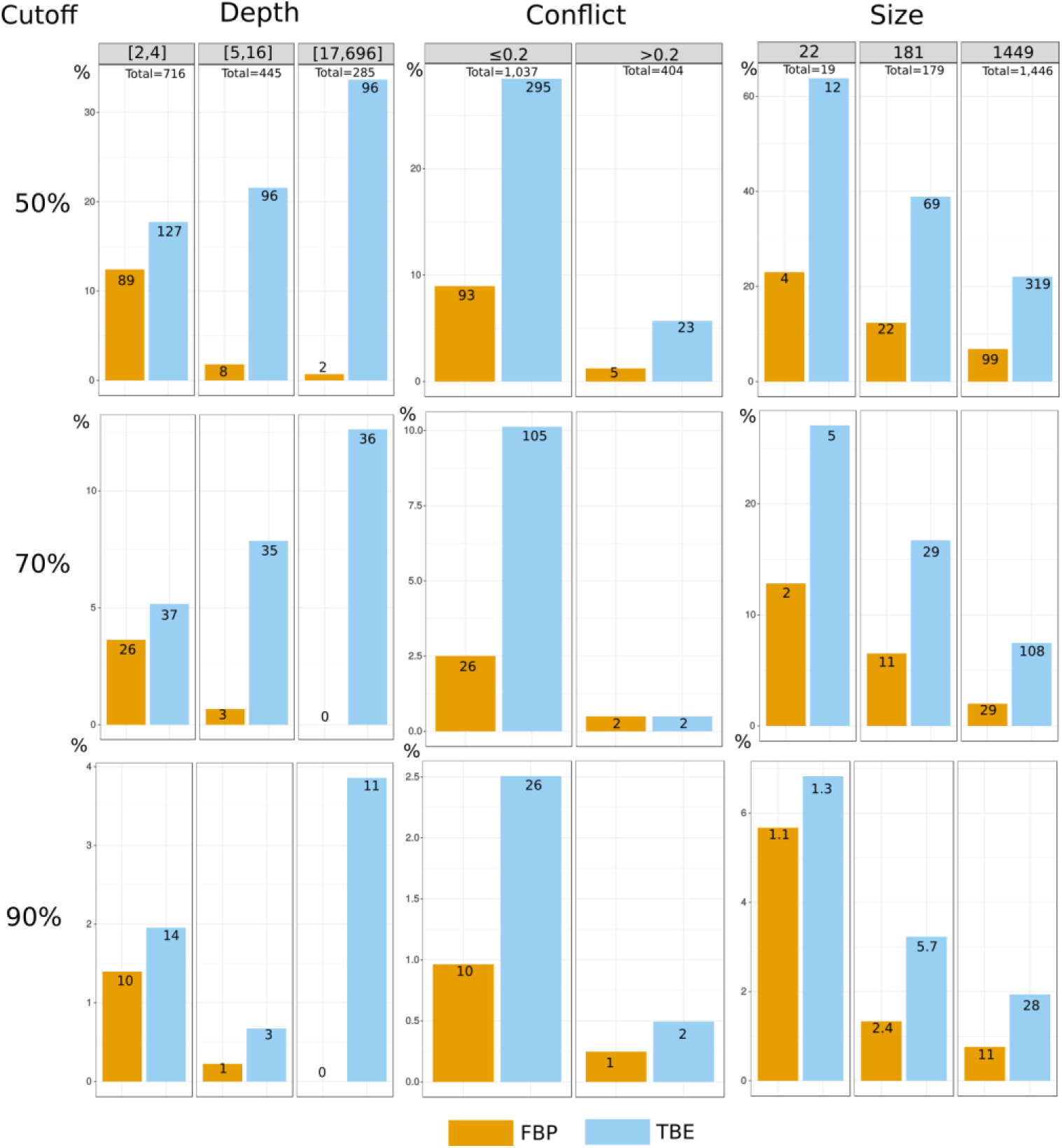
Comparison of FBP and TBE – FastTree – Mammal dataset. Both supports are compared regarding branch depth, quartet conflicts with the NCBI taxonomy, and tree size. Three support cutoffs are used to select the branches: 50%, 70%, and 90% (e.g. 28 branches among 1,446 have TBE ≥ 90% and 11 have FBP ≥ 90%). See notes to Fig. 1 and Fig. 2 for details.

**Fig. S4:**
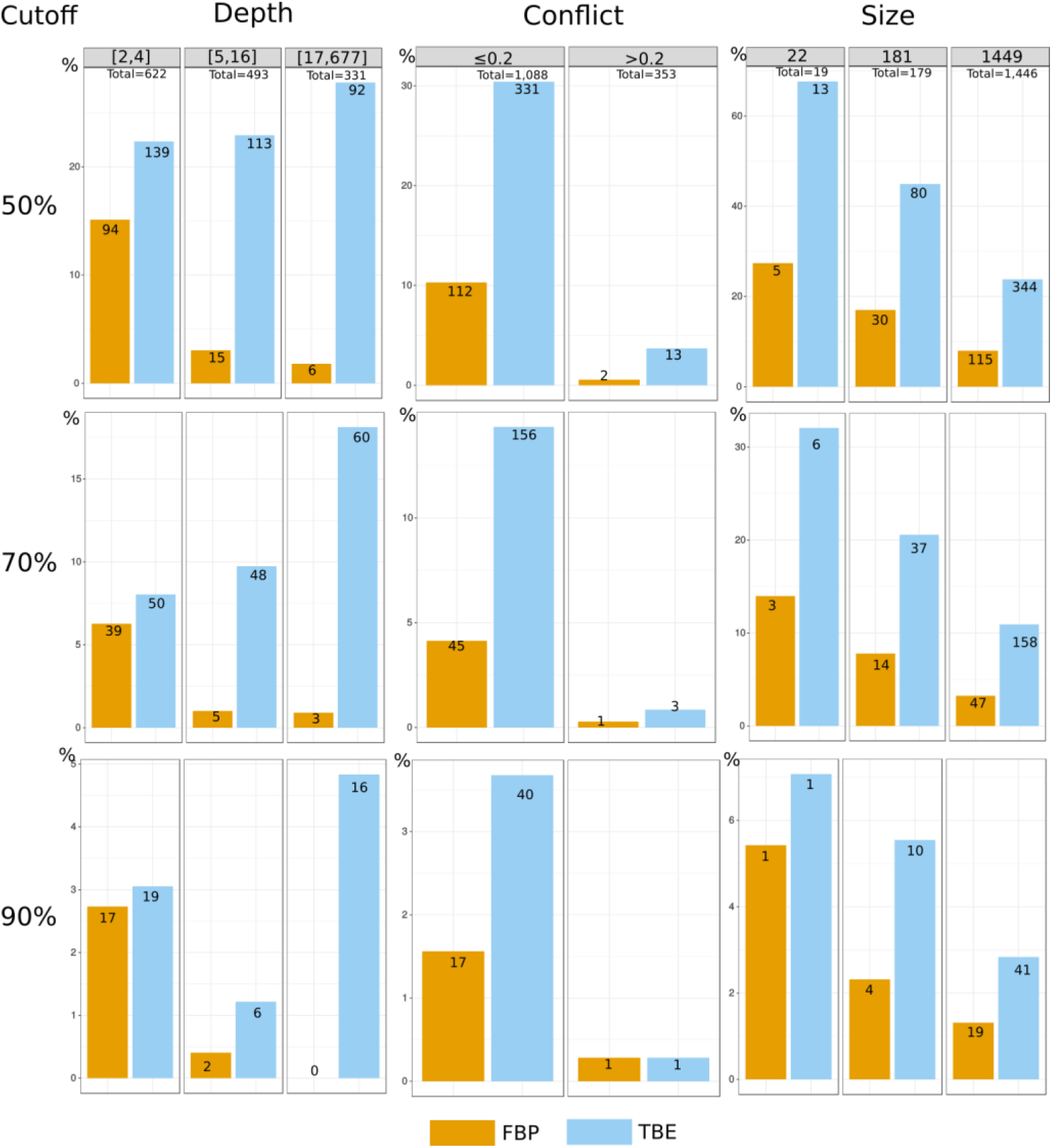
Comparison of FBP and TBE – RAxML with rapid bootstrap - Mammal dataset. See note to Fig. S3.

### Results - Simian clade (Fig. 2, S5)

Fig. 2 displays the FastTree phylogeny for the simian clade; Fig. S5 displays that for RAxML and the supports obtained with rapid bootstrap.

- Both FastTree and RAxML simian trees are well reconstructed and close to the NCBI taxonomy, with less than 2.5% of contradicted quartets (computed using tqDist by comparing the simian subtree from the NCBI taxonomy and pruning from our trees the non-simians). Both clades include all simian sequences (152), but FastTree’s includes two non-simian taxa (*Maxomys rajah* and *Canis adustus)*, while RAxML’s includes *Canis adustus* only. The latter is not a rogue taxon: its sequence is incomplete and very close to the simian sequences for the part being available, and its position is very stable in the bootstrap trees (it is transferred only 2/1000 times when computing the support of the simian clade inferred by FastTree). Oppositely, *Maxomys rajah* is a rogue taxon (656/1000 transfers with FastTree). A number of taxa are rogue-unstable and found in the simian clade in at least 1% of the bootstrap trees: 138 with FastTree and 19 with RAxML. This explains the low FBP support of the simian clade, especially with FastTree.
- Both FBP and TBE supports are again higher with RAxML than with FastTree, although both inferred simian trees have similar, high accuracy (except regarding *Maxomys rajah*). This is the consequence of the higher supports of the rapid bootstrap compared to the standard bootstrap (see above).
- We see the advantage of TBE with both FastTree and RAxML. For example, with RAxML, the simian clade has FBP=50% and TBE=99%, meaning that on average ~1.5 taxa must be transferred to recover the inferred clade, which is the true simian clade, except for the partial *Canis adustus* sequence. Moreover, 64 branches in the simian tree have TBE >70%, versus 19 having FBP >70%, and these additional branches mostly agree with the NCBI taxonomy (average contradiction of ~0.9%).

**Fig. S5:**
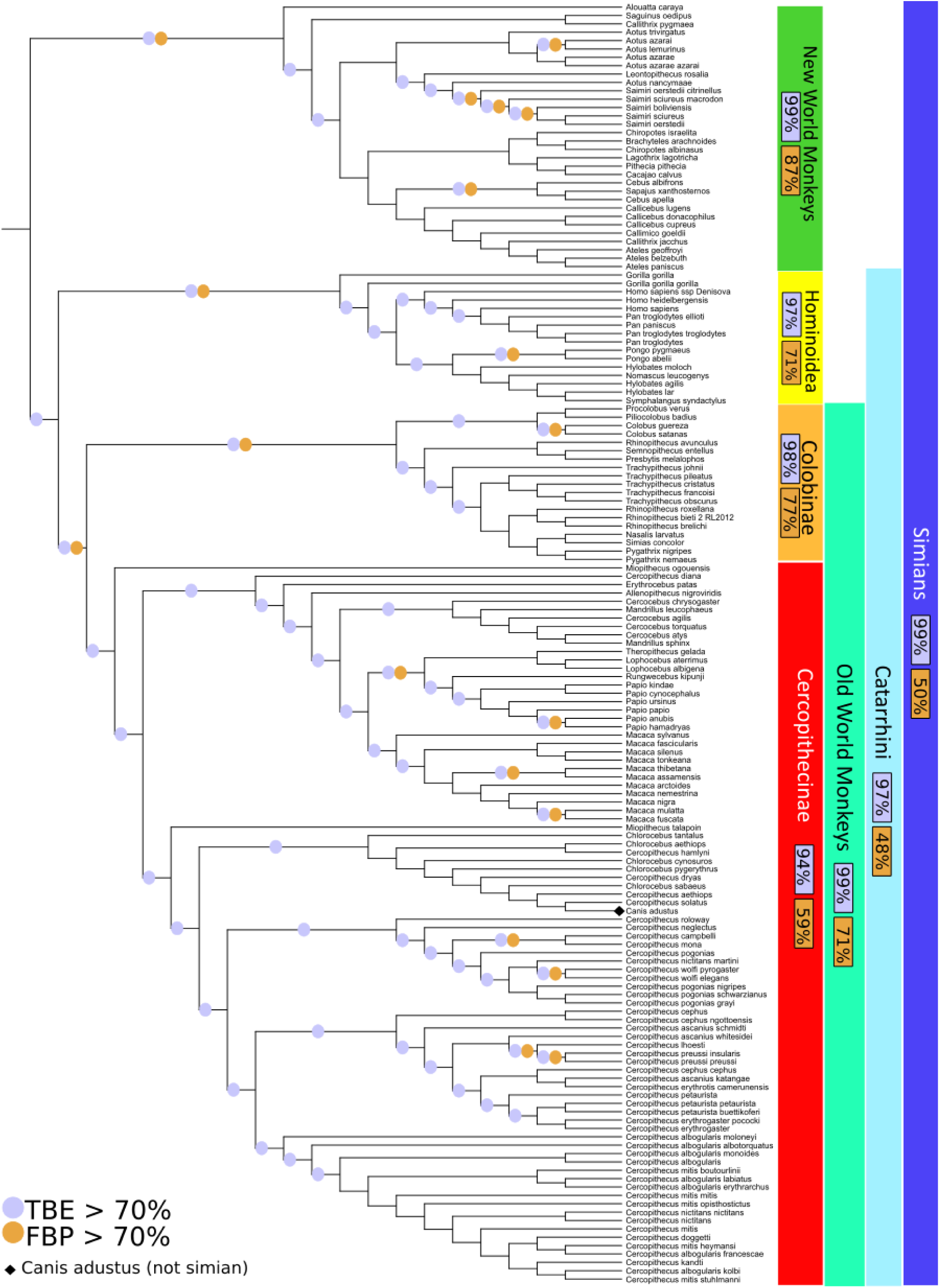
Simian clade – RAxML with rapid bootstrap. See note to Fig. 2.

### Results – Complete phylogeny (Fig. S6)

- The complete phylogeny found by RAxML with rapid bootstrap is shown in Fig. S6. As explained above, this phylogeny is more accurate than that of FastTree, but still relatively poor, especially regarding deep nodes and larger groups. For example, rodents and chiropters are not monophyletic and are distributed in several subtrees.
- FBP does not support any deep branches, except cetaceans (67%), and to some extent simians (50%). FBP also supports some small well-identified clades, namely the monotremes (3 taxa, 92%) and elephantidae (8 taxa, 98%).
- FBP provides strong supports (96% to 100%) for these four groups, but also for five other groups, such as marsupials and insectivores. The latter clade, as inferred by RAxML, contains all insectivores of the NCBI taxonomy, plus one extra taxon. In some sense, both supports are correct: FBP in saying that, strictly speaking, this clade has no support; and TBE in saying that a few taxa are unstable. The advantage of TBE is that this nearly-correct clade is distinguished, as well as the unstable taxa.

**Fig. S6:**
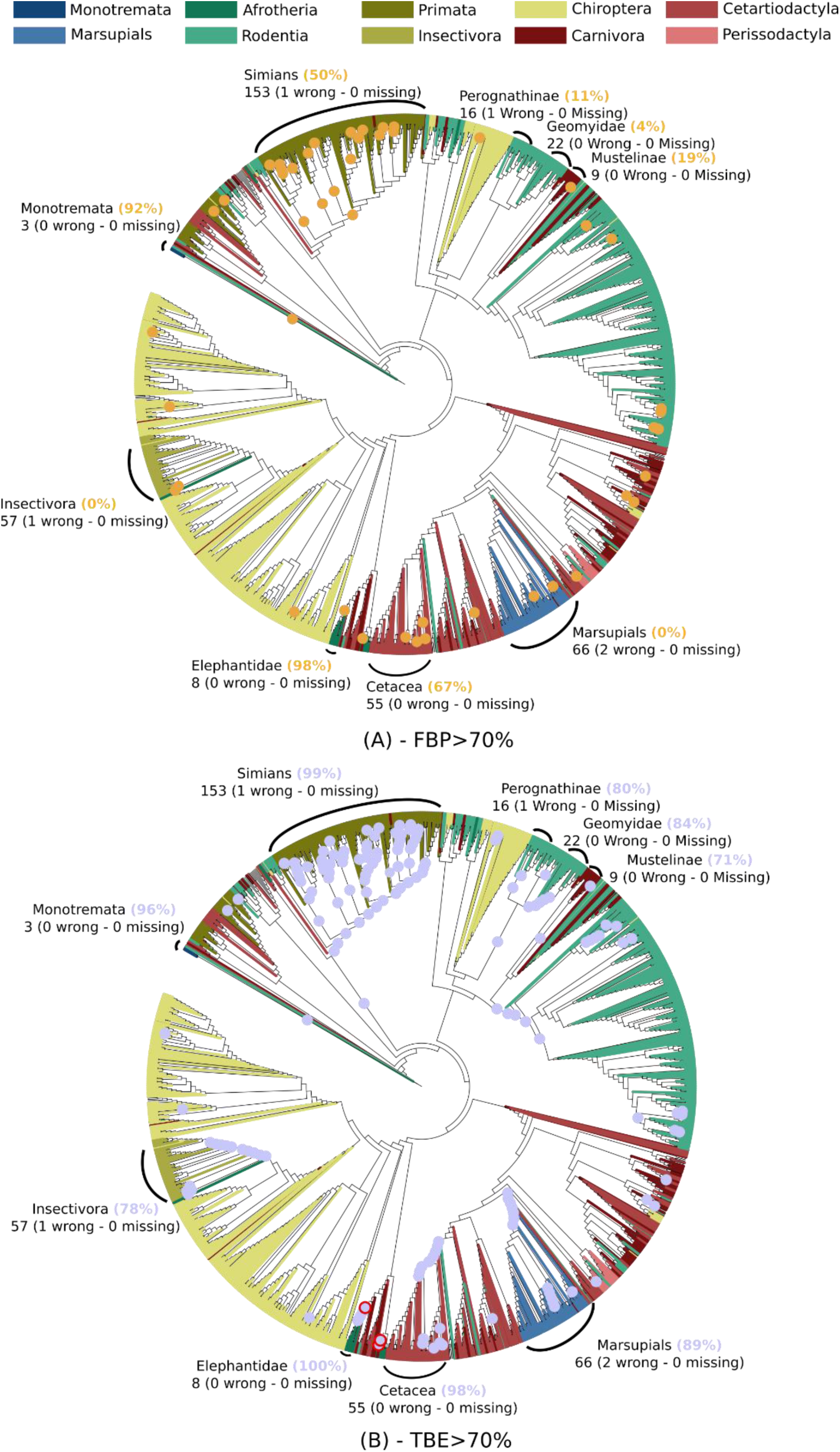
Complete mammal tree – RAxML with rapid bootstrap. A few clades are highlighted, corresponding (almost) exactly to the NCBI taxonomy. For example, all elephantidae taxa are recovered by RAxML in a single clade, containing elephantidae only, while insectivores are included in a clade containing one extra taxon. To select these clades, we minimized the transfer distance with the NCBI taxonomy, in case of ambiguity.

## HIV dataset and analyses (Supporting Text 3)

### Data, alignments, tree building

From the HIV database (https://www.hiv.lanl.gov/content/index) we retrieved *pol* sequences of the nine “pure” subtypes of HIV-1 group M, corresponding to positions 2258-3300 relative to the HXB2 reference strain (access date: September 2014). The “one sequence per patient” option was used, and we randomly selected samples of the over-sampled subtypes (A1, B, C, D, and G), resulting in a final dataset of 9,147 sequences. These sequences are annotated as “pure” (i.e. non-recombinant) in the database, using a fast filtering approach. However, 48 recombinant sequences were still detected using the standalone version of jpHMM ((*30*); version: March 2015; options: -v HIV, with default input and priors). These 48 sequences were kept in the analyses to study the impact of recombinant sequences, as their presence is inevitable in any HIV dataset. jpHMM was also used to annotate the whole set of sequences depending on their subtype or recombinant status.

Sequences were aligned using MAFFT 7.0 ((*37*); default parameters) along with the HXB2 reference strain. Codon positions associated with major drug resistance mutations were removed prior to tree inference, resulting in an alignment of 1,043 DNA sites (R source code available at https://github.com/olli0601/big.phylo). This alignment was sub-sampled to study the impact of tree size. We randomly drew 16 samples with 1/16th of the sequences (i.e. 571), and 256 samples with 1/256th of the sequences (i.e. 35). Then, we generated 1,000 bootstrap alignments for the full alignment and each of the 272 sub-sampled alignments, by drawing sites with replacement.

We used FastTree ((*28*); options: -nopr -nosupport -gtr -nt -gamma) to infer trees from each of these reference (1+16+256=273) and bootstrap (273,000) alignments. The FBP and TBE supports for the 273 reference trees were computed using Booster (command-line C version). All trees were drawn using iTOL (http://itol.embl.de/) and are available on the Booster-workflows GitHub repository, along with sequence alignments. The instability score was computed considering the reference branches with TBE >70% (the signal becomes noisy when incorporating branches with lower supports in the calculation, as these branches may be erroneous and thus noninformative about taxon stability). For every taxon, the instability score is equal to the average number of times it has to be transferred to recover these branches from the bootstrap trees, divided by the number of these branches.

The most representative clades for each of the subtypes in the reference trees (Fig. 1, Fig. S8) were obtained by minimizing the transfer distance. For example, in Fig. 1 with the full dataset, we obtained a clade very close to subtype B, with 3,559 taxa, 2 wrong taxa (i.e. non-B), and all B taxa included (i.e. 3,557, 0 missing).

A similar approach was used for the regional variants of subtype C, which is responsible for approximately 50% of the HIV-1 infections in the world. Three monophyletic variants of subtype C have been identified by phylogenetic analysis in East Africa (*38*), South America (*39*), and India (*40*). Moreover, the South American epidemics was shown to originate from the East African cluster (*38*). To identify these variants in the inferred trees (Fig. 1, Fig. S8) we again used the transfer distance. Following (*38-40*), we extracted three groups of C sequences from the whole dataset, based on their geographic origins: East Africa (EA 440 sequences = Burundi 288 + Djibouti 1 + Ethiopia 9 + Kenya 41 + Somalia 1 + Sudan 11 + Tanzania 78 + Uganda 11); India (IND 154 sequences = India 133 + Nepal 13 + Myanmar 8); and South America (SA 14 sequences = Brazil 12 + Uruguay 1 + Argentina 1). We then searched for the tree clades being closer to these three sets of sequences. The SA sequences were not accounted for in transfer distance computations when searching for the EA clade, as they originate from EA. Moreover, we checked that no neighboring, nearly-optimal clade was supported by FBP. In all three cases, we found clades closely related to the sequence sets. As expected, the SA clade was included in the EA clade. The features of these clades are displayed in Fig. 1 and S8. The fractions correspond to the number of studied sequences included in these clades, versus the total number of such sequences in the whole dataset (e.g. 360 EA sequences in the EA clade in Fig. 1, among 440 in the whole tree). The “wrong” sequences were expected in most cases. For example, the IND clade (167 sequences, 143 from IND among 154 in the whole tree) contains 19 sequences from China corresponding to the spread in Asia via heroin trafficking routes (*40*).

### Results – Overall comparison of FBP and TBE (Fig. 1, S7)

- Results are mostly similar to those observed with the mammal dataset. Again, we see a major impact of the depth on FBP supports: with the full dataset, less than 1% of the deep (>16) branches have FBP support larger than 70%, whereas this percentage is higher than 20% with TBE.
- The impact of tree size is less pronounced. The fraction of supported branches decreases when the tree size increases from 35 to 571 taxa, but is analogous between 571 and 9,147 taxa. Moreover, the gap between FBP and TBE remains similar, likely due to the very large number of cherries and small clades (5,478 among 9,144 with the full dataset), where TBE and FBP are nearly equivalent.
- Regarding the cutoff, 70% again appears as a good compromise for TBE, though there is no way to evaluate the fraction of supported branches that are actually erroneous. As with mammals, the interpretability of TBE will be a major asset for choosing the support level depending on the phylogenetic question being addressed. Here, as recombinant sequences are inevitable, lower supports than with mammals will likely be acceptable.

**Fig. S7:**
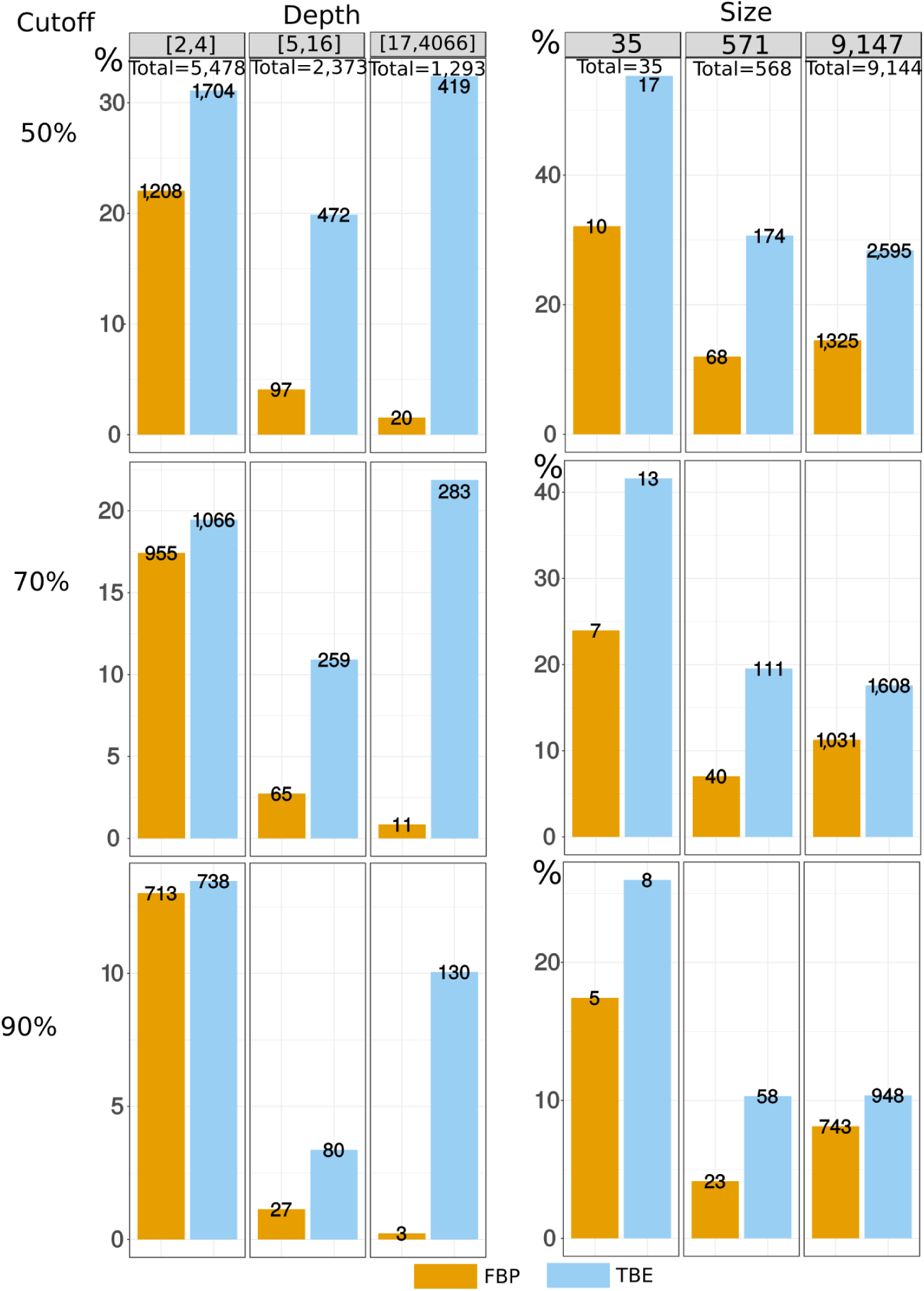
Comparison of FBP and TBE – HIV-1M *pol* dataset. See notes to Fig. 1-2, S3-S4.

### Results – Medium-sized datasets (Fig. S8)

- Results with 571 taxa are shown in Fig. S8. As the taxa were randomly drawn from the full dataset, the supports and findings show some fluctuations, and we display the trees obtained with two of the medium-sized datasets. Rare subtypes (H, J, K) are absent, and the subtype clades are almost perfectly recovered (only 1 wrong taxon in A clade for both trees).
- FBP supports are higher than with the full dataset (e.g. 58% and 99% for subtype B, versus 3% in Fig. 1). However, some subtype clades have moderate FBP (e.g. D), though the clade matches the subtype perfectly. With TBE, all subtype supports are higher than 95%.
- The deep branching is the same for all (full, medium) datasets and identical to (*19*), but not supported by FBP, while TBE is larger than 70% for every branch (or path in Fig. 1).

**Fig. S8:**
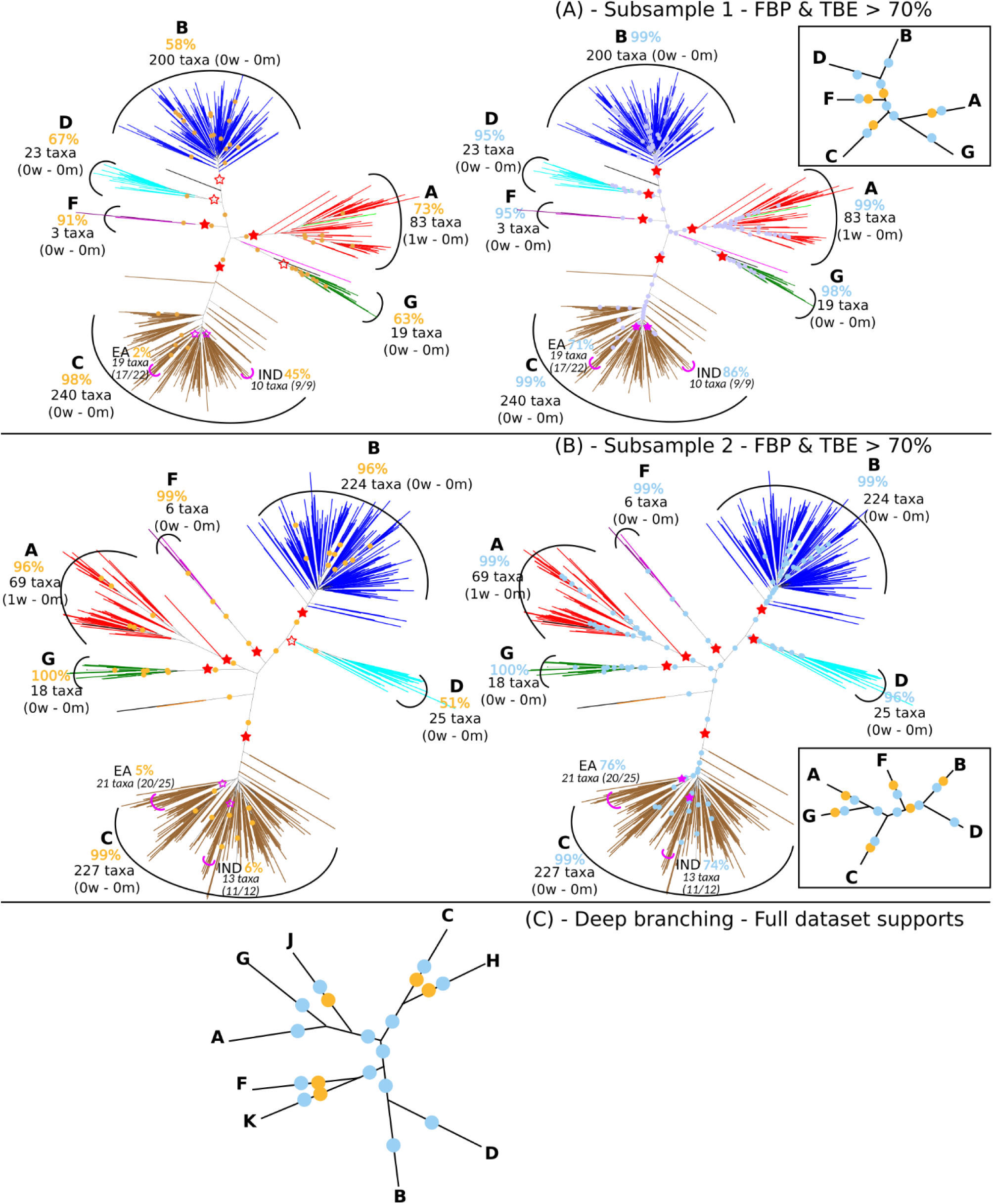
Results with medium-sized datasets and subtype deep branching. (A) and (B): branches with FBP>70%: yellow dots; branches with TBE>70%: blue dots; subtype clades: red stars, filled if support >70% (see notes to Fig. 1 for further details). (C) Deep branching of the subtypes (*19*) and supports obtained on the full data set (Fig. 1).

### Results – Instability score (Fig. S9)

- We see a clear difference between the distributions of the instability score for the recombinant and non-recombinant sequences, meaning that the approach could be used to detect or confirm the recombinant status of sequences.

**Fig. S9:**
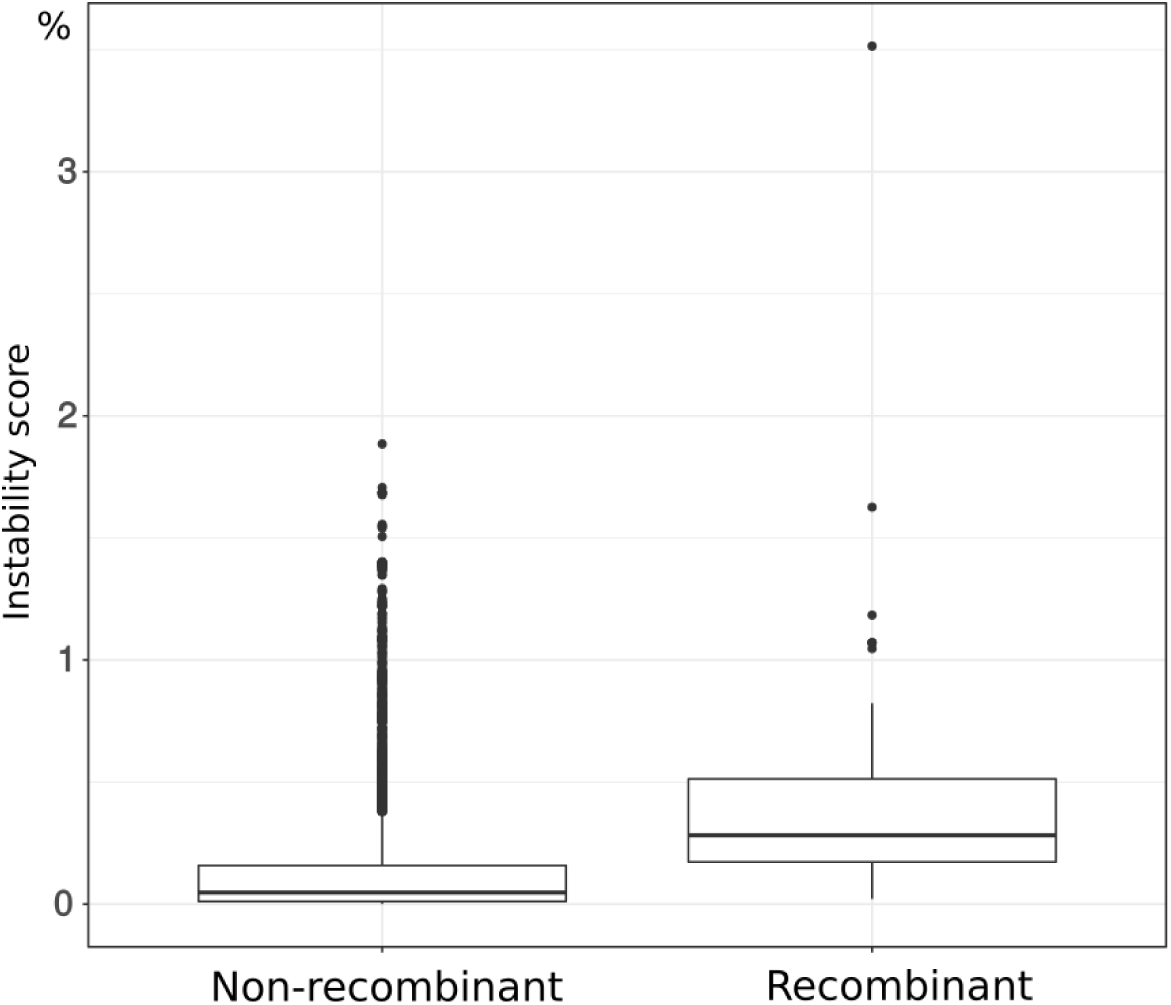
Distribution of the instability score among recombinant and non-recombinant sequences. See text for details.

## Simulated data and analyses

The aim of these simulation experiments was to check that the results observed with the mammal and HIV-1 datasets are reproducible and quantifiable when the simulation conditions and correct tree are known, notably regarding the support of poor branches and the ability to detect rogue taxa. Simulated data mimicked the mammal dataset. We used the tree inferred by PhyML ((*41*); options: -b 0 -m WAG -a e –t e -o tlr -d aa) from the full COI-5P protein alignment with 1,449 taxa. Protein sequences with length 300 (plus gaps) were evolved along this tree using INDELible (*42*), which was launched with options and parameter values derived from the PhyML analysis, and similar to the experiments conducted in (*24*) to assess the accuracy of rogue-taxon detection:

- Substitution model: WAG;
- Amino-acid frequencies estimated from the COI-5P alignment;
- Rates across-sites model: 4 gamma categories with ‘alpha’ =0.441 and no invariant sites;
- Indel model: ‘power law’, ‘parameter’ =1.5, ‘indel max size’ =5, and ‘indel rate’ = 0.02.

We so obtained a first easy, “non-noisy” MSA. Noise was added to this MSA to mimic rogue taxa and homoplasy. We shuffled the amino acids vertically for 50% of the sites (MSA columns), thus making these sites homoplasic. For 5% of the sequences (MSA rows), 25% additional sites were shuffled vertically, thus making these sequences unstable and “rogue”, as they contain half of the phylogenetic signal compared to the 95% others. Both noisy and non-noisy MSAs were used to compare FBP and TBE. To measure the effect of tree size, both MSAs were sampled to obtain 8 MSAs with 181 sequences (~1/8 of the full sequence set) and 64 MSAs with 22 sequences (~1/64 of the full sequence set). For each of these reference MSAs we sampled with replacement 1,000 pseudo-alignments to compare the two bootstrap methods. All trees were inferred using FastTree (options: -nopr -nosupport -wag -gamma). Just as with the mammal dataset, we computed, for each of the branches in the reference trees, the percentage of quartet-based conflicts with the correct (PhyML) tree used to generate the data. We also computed the instability score of all taxa in the complete noisy MSA, using only the branches with TBE>70%.

Results (Fig. S10-S12) are fully congruent with those already shown with real datasets. TBE supports more deep branches than FBP, especially with noisy data. The effect of tree size is also more visible with noisy MSA, and the percentage of conflictual branches being supported is extremely low and similar for FBP and TBE. Regarding the support cutoff, 50% seems to be acceptable, as neither FBP nor TBE support highly contradicted branches. But this could be due to the low level of contradiction, compared to real datasets (85 branches with contradiction >20% in Fig. S10-S11, versus ~400 with the mammal dataset in Fig. S3-S4). Moreover, TBE again appears to be useful for detecting and confirming rogue taxa (Fig. S12).

**Fig. S10:**
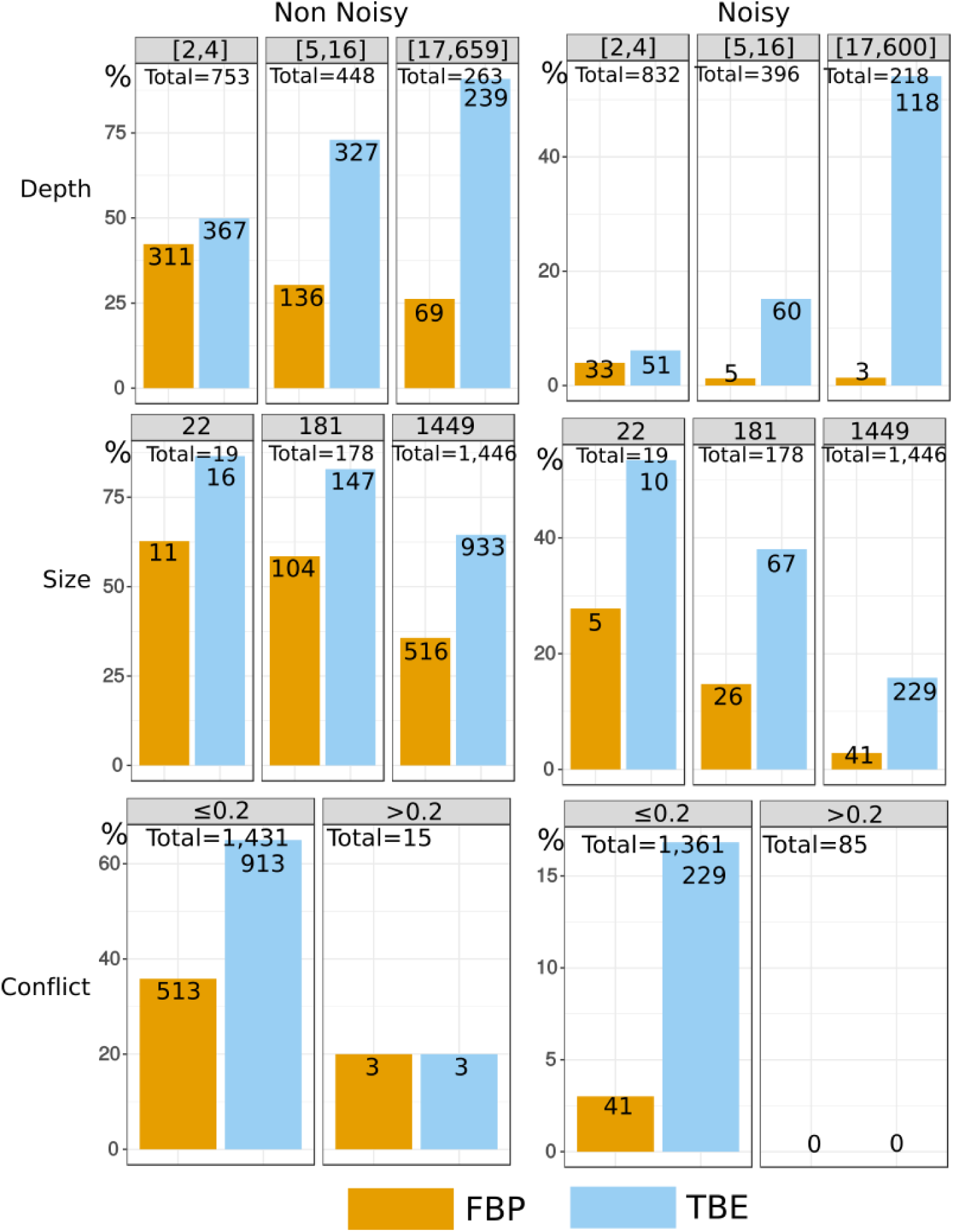
Results with simulated data. These graphics display the distribution of branches with FPB/TBE support >70%, for both the non-noisy and noisy MSAs. See text and note to Fig. S3 for details.

**Fig. S11:**
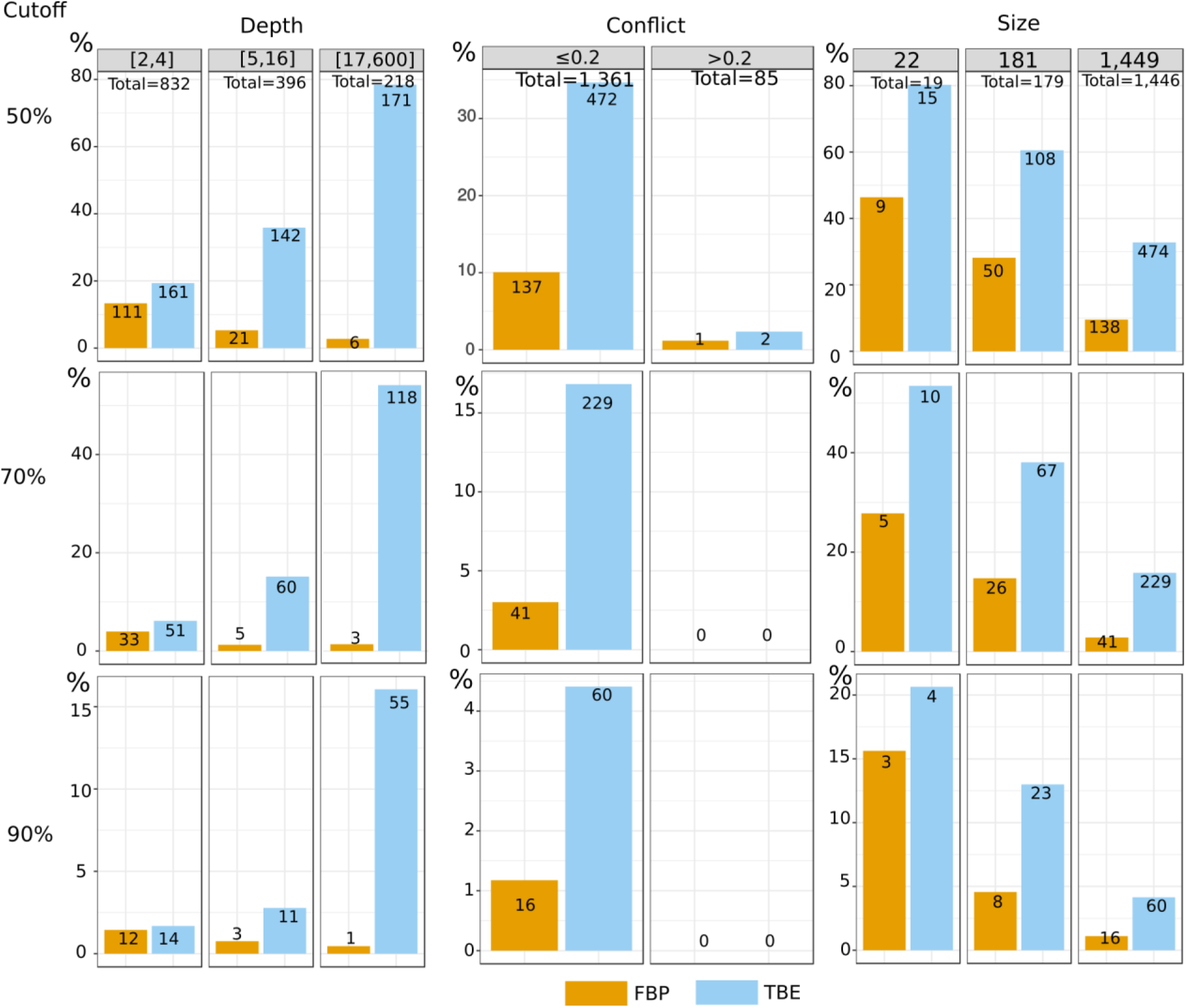
Results with simulated data - noisy condition. Comparison of FBP and TBE regarding branch depth, quartet conflicts, and tree size, at different support cutoffs. See note to Sup Fig. S3 for details.

**Fig. S12:**
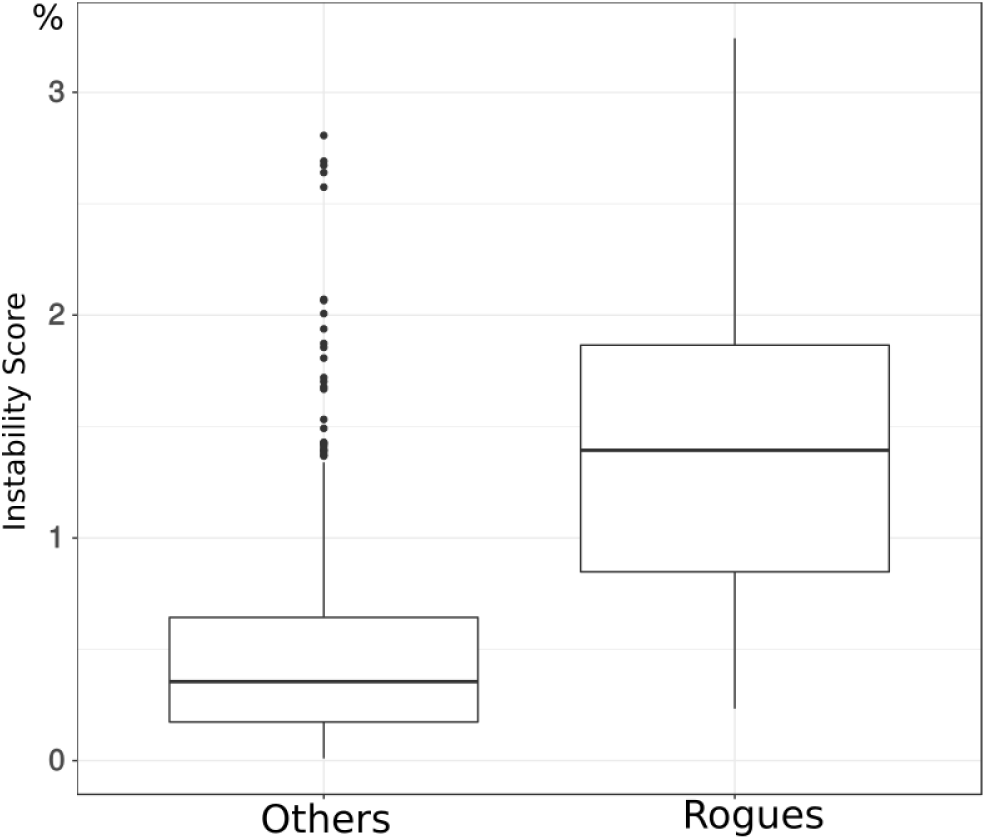
Distribution of the instability score with simulated data - noisy condition. See text for details.

